# Partial-Width Injuries of the Rat Rotator Cuff Heal with Fibrosis

**DOI:** 10.1101/268920

**Authors:** Elisabeth A. Lemmon, Ryan C. Locke, Adrianna K. Szostek, Megan L. Killian

## Abstract

**Purpose:** The purpose of this study was to identify the healing outcomes following a partial-width, full-thickness injury to the rotator cuff tendon-bone attachment and establish if the adult attachment can regenerate the morphology of the healthy attachment.

**Hypothesis:** We hypothesized that a partial-width injury to the attachment would heal via fibrosis and bone remodeling, resulting in increased cellularity and extracellular matrix deposition, reduced bone volume, osteoclast presence and decreased collagen organization compared to shams.

**Materials and Methods:** A biopsy punch was used to create a partial-width injury at the center one-third of the rat infraspinatus attachment, and the contralateral limb underwent a sham operation. Rats were sacrificed at 3- and 8-weeks after injury for analyses. Analyses performed at each time-point included cellularity (Hematoxylin & Eosin), ECM deposition (Masson’s Trichrome), bone volume (micro-computed tomography; microCT), osteoclast activity (Tartrate Resistant Acid Phosphatase; TRAP), and collagen fibril organization (Picrosirius Red). Injured and sham shoulders were compared at both 3- and 8-weeks using paired, two-way ANOVAs with repeated measures and Sidak’s correction for multiple comparisons.

**Results:** Cellularity and ECM deposition increased at both 3- and 8-weeks compared to sham contralateral attachments. Bone volume decreased and osteoclast presence increased at both 3- and 8-weeks compared to sham contralateral limbs. Collagen fibril organization was reduced at 3-weeks after injury compared to 3-week sham attachments.

**Conclusions:** These findings suggest that a partial-width injury to the rotator cuff attachment does not fully regenerate the native structure of the healthy attachment. The injury model healed via scar-like fibrosis and did not propagate into a full-width tear after 8-weeks of healing.

## INTRODUCTION

Rotator cuff tears are a common orthopedic injury, with over 75,000 surgical repairs performed annually (1). Depending on the location and severity of the tear, clinical recommendations for surgery and physical therapy vary substantially (2, 3). For partial-width tears, conservative treatments (e.g., physical therapy) are encouraged before surgical intervention (3–5). The success of rotator cuff repair depends on the health of the cuff at time of repair (6–10) as well as the restoration of the morphology and strength of the attachment (11–16). Following full-width tears, repair of the torn attachment rarely results in full regeneration of the native morphology and structure of the attachment (6, 14, 17–19). In small animal models of cuff healing, a fibrous scar tissue, characterized by reduced collagen organization, increased vascularization, and increased cell density, are often observed (14, 18, 19). Loss of the structural integrity of the attachment is considered one of the primary causes of high rates of re-injury after rotator cuff repair (6, 19). However, animal models used to study rotator cuff healing have primarily relied on full-width injuries that require surgical reattachment of cuff tendons to their bony footprint for structural reintegration. Healing of cuff tendons during early stages of tear propagation, such as in the case of partial-width injuries, have been investigated *in vivo* using animal models (19–22). However, the healing of partial-width injuries focused at the attachment, without surgical repair and augmentation, have only recently been explored (21).

In the present study, we aimed to develop and validate a new model of rotator cuff injury to investigate the natural healing process of the attachment. Using a rat model of partial-width, full-thickness injury at the attachment, we hypothesized that, although the attachment would remain partially intact, it would not heal with the same structural quality as an intact, uninjured attachment. We tested this hypothesis by assessing the fibrotic response, extra-cellular matrix deposition, bone remodeling, and collagen organization of the healing attachment.

## MATERIALS AND METHODS

### Animal Model

Adult Sprague Dawley rats (N = 8 females, N = 8 males for *in vivo* healing; ~200-250g) and adult Long Evans rats (N=20 female dams, for *ex vivo* validation at time zero) were used in accordance with the University of Delaware Institutional Animal Care and Use Committee approval. Rats underwent a surgical procedure under anesthesia (isoflurane carried by 1% oxygen) to model a partial-width rotator cuff injury at the center of the IS attachment (Figure 1A–C) (23). The IS tendon was exposed and the forearm was internally rotated. A 0.3mm diameter biopsy punch (Robbins, Chatham, NJ, USA) was placed in the center 1/3 of the tendon width spanning the attachment/bone, and the punch permeated the fibrocartilage, tendon, and cortical bone. The injured shoulder was randomized between rats, and the contralateral shoulder underwent a sham operation to mimic the procedure without the biopsy punch permeation. The surgical site was closed using 5-0 Vicryl suture (Ethicon Inc., Somerville, New Jersey), and rats were given bupivacaine hydrochloride (0.05 mg/kg) as analgesia. Rats were separated into two groups: 3-week healing (N=8; 4 females and 4 males) and 8-week healing (N=8, 4 females and 4 males). Rats were sacrificed with carbon dioxide asphyxiation and thoracotomy. The 3-and 8-week time points were chosen to evaluate the proliferation (3-week) and remodeling (8-week) phases of tendon-bone healing.

**Figure 1.**
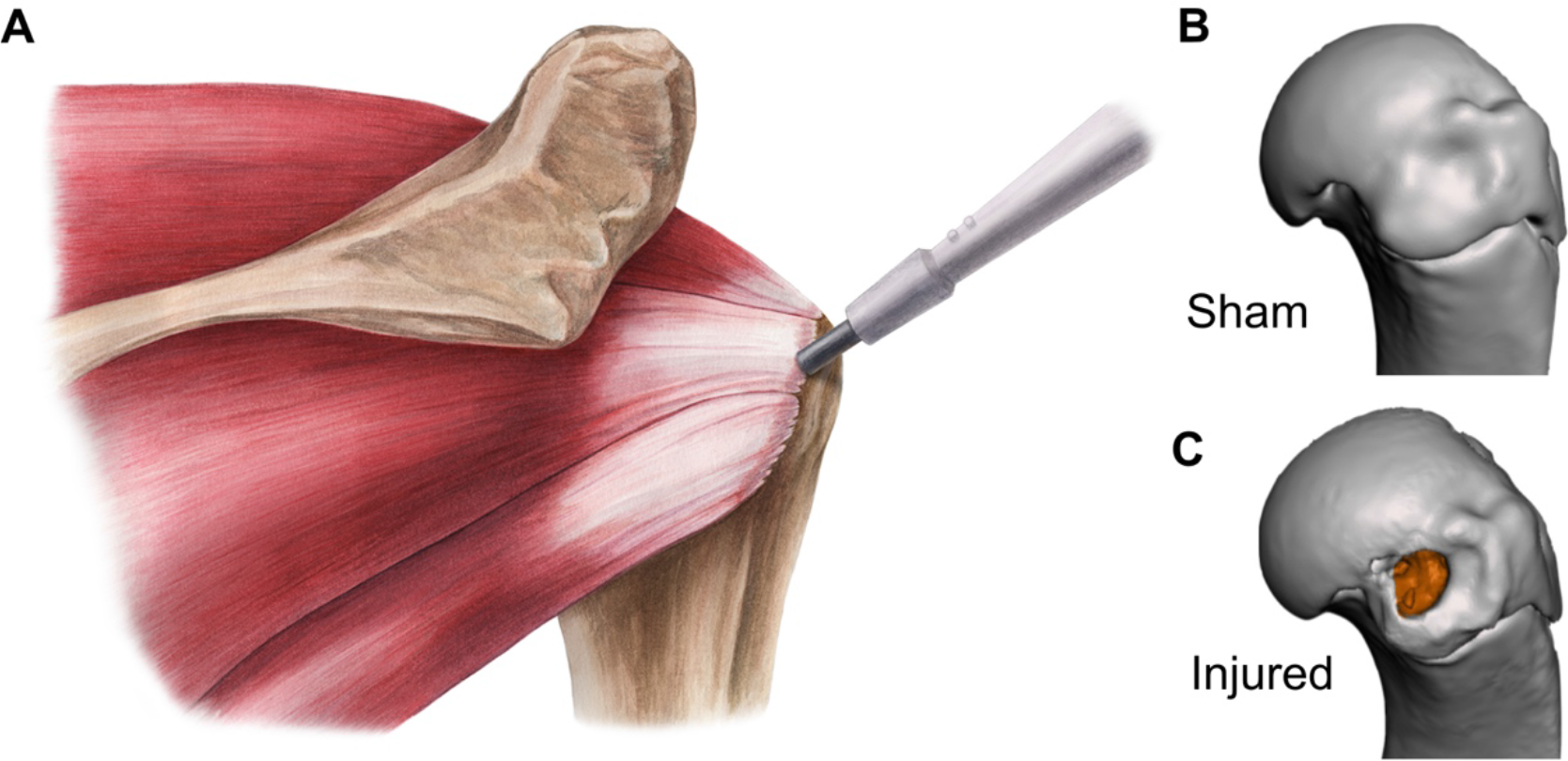
Rotator cuff infraspinatus injury model and microCT reconstructions of sham and injured groups. (A) Illustration of the human rotator cuff muscle groups. In our animal model, a 0.3mm circular injury was made at the center of the infraspinatus attachment. MicroCT renders of (B) sham and (C) injured humeral heads. The orange region is the injury location.

### Biomechanics

Validation of the model was evaluated *ex vivo* using biomechanical testing (on N = 18 rats). Adult, female Long Evans rats, used as breeders in an unrelated study (Department of Psychology and Brain Sciences), were sacrificed after weaning of their first-born litters. Dams were not treated with any pharmacological drugs in previous studies. Superficial (through soft tissue/fibrocartilage) and deep (permeating the cortical bone) injury defects (N = 8-10 shoulders each) were made evaluating the biomechanical properties of the defect at time zero and establish whether a superficial or deep defect result in comparable biomechanical outcomes. Rats were randomized as to which limb received the defect, and the contralateral limb underwent a sham operation to mimic the procedure, similar to *in vivo* approaches. After performing the punch, the IS tendon-to-bone attachments were dissected from the surrounding musculature and bone. The IS muscle belly was detached from the IS fossa of the scapula and the IS tendon attachment at the proximal humerus was left intact for mechanical testing. Uniaxial tensile tests were performed (Instron 5943, Norwood, MA) and the cross-sectional area of the attachments were measured using microCT. To prevent failure at the growth plate during testing, a hole was drilled in the humeral diaphysis and then steel wire was passed through the hole and wrapped posterior to anterior around the humeral head. Tendons were tested in a PBS bath using custom fixtures to ensure uniaxial loading. A 0.01N tare load was applied and the test configuration was imaged at the start of the test to measure gauge length of the sample, followed by five preconditioning cycles to 5% strain at a rate of 0.2%/sec. Following preconditioning, samples were held for 30s at initial gauge length and then loaded to failure at 0.2%/sec. Load/displacement data were recorded throughout the test (preconditioning and load-to-failure) and data were converted to stress/strain data based on the initial cross-sectional area from microCT (stress) and the gauge length from the start of the test (strain). Stiffness (N/mm) and ultimate load (N) were calculated from load/displacement curves, and Young’s modulus (MPa) and ultimate stress (MPa) were calculated from stress/strain curves, with stiffness and Young’s modulus as the slope of the respective curves in the linear region and ultimate load/stress as the maximum load/stress of the respective curves. Data were calculated using MATLAB (MathWorks, Natick, MA, USA).

### Micro-Computed Tomography

We validated our injury model via micro-computed tomography (microCT) *ex vivo* to ensure the correct location of the punch at the attachment at time of injury (Supplemental Figure 1A-F). The shoulder complexes of female Long Evans rats (N = 2 for each group; previously sacrificed) were used within two hours of sacrifice and unilateral shoulder complexes were exposed to perform partial-width injuries with the biopsy punch at superficial (through soft tissue/fibrocartilage) and deep (permeating the cortical bone) intervals. For both validation and *in vivo* healing studies at 3-and 8-week healing, the shoulders were carefully removed to isolate the IS tendons and corresponding muscles. Humeri were cut at the mid diaphysis distal to the deltoid tuberosity using bone shears. Shoulders were fixed in their anatomical position in 4% paraformaldehyde (N = 2 each for control, superficial, and deep injuries for validation studies; N = 8 each for 3- and 8-week healing time points). Shoulders were scanned in air using microCT (Scanco µCT35; 20-µm voxel size, 45kV, cone beam, 177µA, 800-msec integration time). The tendon-bone attachments were analyzed for structural quality based on 3-dimensional reconstructions produced in OsiriX DICOM Viewer (v8.0.2., Pixmeo, Switzerland) and quantitatively assessed using Scanco software. Bone morphometric properties were quantified for the 1) entire humeral head and 2) the injury region using total volume (TV), bone volume (BV), and bone volume/total volume ratio (BV/TV) (Supplemental Figure 2A-B). The humeral head measurements included the humeral head proximal to the growth plate, trabecular bone, and cortical bone. The injury region measurements comprised the IS tendon-bone region proximal to the growth plate, including both the trabecular and cortical bone (Scanco, Switzerland).

### Histology

After microCT, shoulders were decalcified in 14% EDTA (Sigma-Aldrich, St. Louis, MO, USA) and processed for paraffin sectioning. Attachments were sectioned at 6µm thickness and stained using Hematoxylin & Eosin (H&E) for cellular density, Picrosirius Red for collagen organization, Masson’s Trichrome for fibrosis and extra-cellular matrix (ECM) localization, and Tartrate Resistant Acid Phosphatase (TRAP) for osteoclast staining. Stained sections were imaged using an epifluorescent microscope (Axio.Observer. Z1, Carl Zeiss, Thornwood, NY). Cell density within the attachment was measured using a custom MATLAB code on a similarly sized region of interest between samples (MATLAB, Supplemental Figure 3A-B’) (24). Sections stained with Picrosirius Red were imaged using circular polarized light microscopy, and the deviation of aligned collagen fibrils was evaluated using a custom MATLAB code (Supplemental Figure 4A-B) (25). TRAP-stained sections were analyzed using Osteomeasure software (Ostemetrics, Decatur, GA, USA) to quantify osteoclast surface area at the injury site (Oc.S.) relative to the total bone surface area (B.S.).

### Statistical Analyses

All statistical comparisons were performed using Prism (v7.0, Graphpad, La Jolla, California, USA). To determine if sex differences influenced our microCT outcomes, an ordinary three-way ANOVA was performed (injury, sex, and time point) with a Tukey multiple comparisons test. If sex had very little impact, we consolidated the data for comparisons using a two-way ANOVA. Within animal comparisons for humeral head BV/TV, injury region BV/TV, cell density, Picrosirius Red circular standard deviations, and Oc.S./B.S were analyzed using two-way ANOVAs (3wk vs. 8wk) with repeated measures (sham vs. injured) and Sidak’s correction for multiple comparisons. All quantitative data are presented as mean ± 95% confidence intervals (CI).

## RESULTS

Based on validation biomechanical results, we performed the deep injury *in vivo* instead of the superficial injury, as the deep injury resulted in significant decreases in stiffness, Young’s modulus, ultimate load, and ultimate stress compared to the uninjured control (Supplemental Figure 1). The superficial injury did not lead to significant differences in biomechanical properties compared to uninjured controls (Supplemental Figure 1).

### Gross Observations

At the time of dissection for both the 3-and 8-week time points, the injured attachments were intact and none of the injuries had developed into full-width tears. No obvious morphometric differences were observed between the injured and sham muscle bellies or humeral heads. Gross examination of the healed injury site indicated increased fibrosis of the fascia surrounding IS tendon-bone attachments.

### Micro-Computed Tomography

Sex did not have a significant impact on the bone morphometry outcomes, and therefore we consolidated sex as a variable for two-way ANOVA (factors: side and time point with repeated measures of side). Similar to our validation results, which showed that the deep injury permeated the cortical bone at time zero, the *in vivo* deep injury permeated the cortical and trabecular bone at the IS attachment, which did not return to its normal morphology by 8-weeks post injury (Figure 2A–F). As visualized in microCT reconstructions, 3-week injured attachments showed reduced amount of bone at the defect site; by 8-weeks post-injury, the defect was mineralized, although this remodeling was incomplete (Figure 2C & F). Quantitatively, humeral head BV/TV was significantly lower at both 3- and 8-weeks for the injured groups compared to the sham groups (combined male and female groups, Figure 2G). In addition, BV/TV increased with age of the rat for both male and female rats, regardless of injury group (Figure 2G).

**Figure 2.**
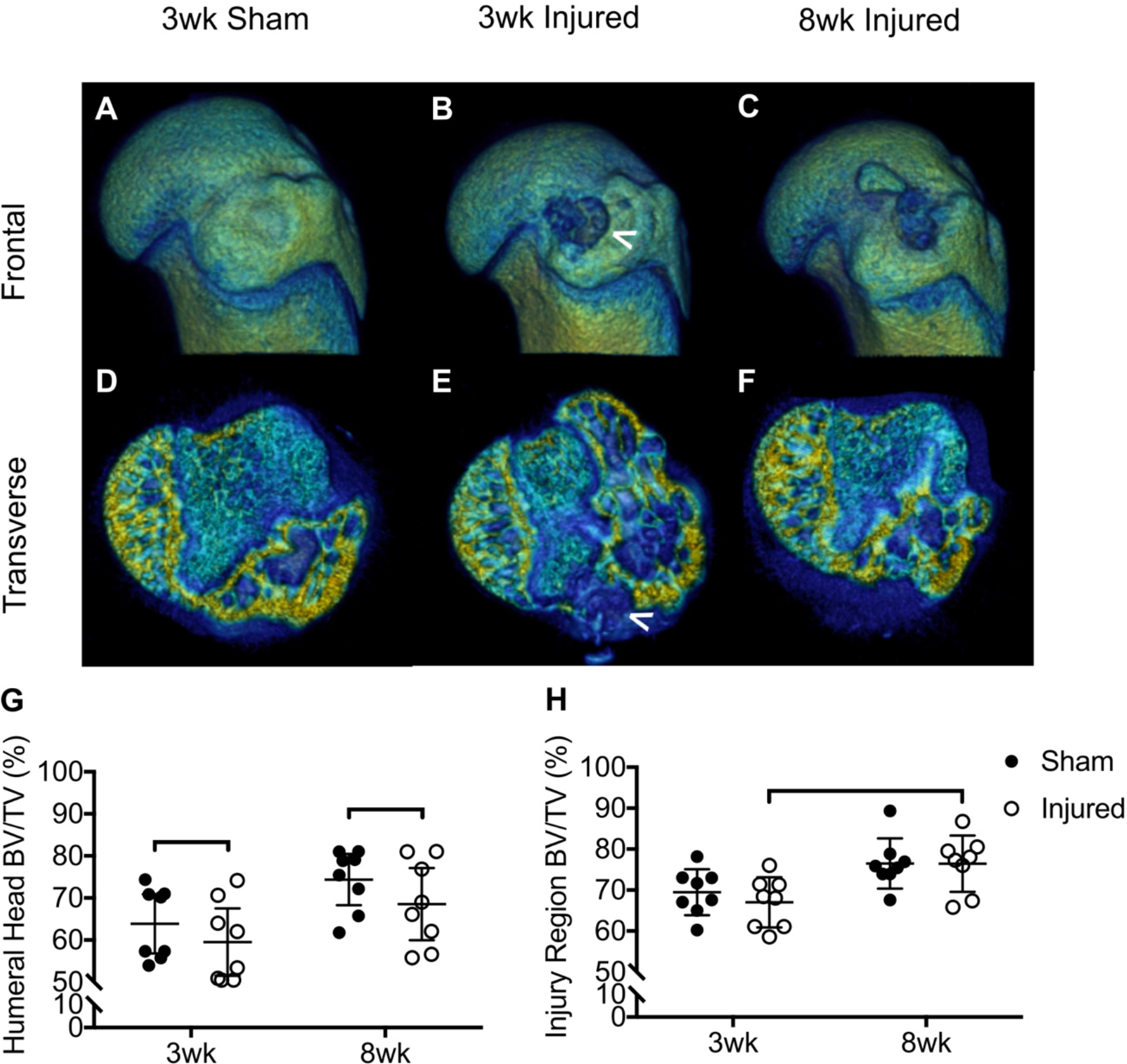
Humeral head BV/TV was significantly reduced in injured attachments compared to the sham forearms for both time points. (A-F) MicroCT reconstructions of sham and injured forearms at 3- and 8-weeks. (A-C) Frontal plane views of microCT reconstructions showing the IS ridge of the humeral head for (A) 3wk sham, (B) 3wk injured, and (C) 8wk injured attachments. (D-F) Transverse cut-plane (D) 3wk sham, (E) 3wk injured, and (F) 8wk injured. White arrowheads: Cortical bone injury in the injury region. (G) Humeral head BV/TV, (H) Injury region BV/TV for 3wk control, 3wk injured, 8wk control, and 8wk injured attachments. Bars indicate significant differences between groups (p < 0.05, mean ± 95% CI).

The injury region had significantly reduced BV/TV in at 3-week post-injury compared to the 8-week injured group (Figure 2H). There were no significant differences in injury region BV/TV between the 3-and 8-week sham (Figure 2H). Measurements for both the humeral head total volume of the attachments as well as injury region total volume were not significantly different across all groups (Supplemental Figure 2C-D).

### Histology

Qualitatively, the injured attachments showed increased ECM deposition and decreased collagen organization, as seen using histological imaging. At 3-week post-injury, increased ECM deposition (shown in red+ staining) was apparent compared to sham attachments (Figure 3B–B’). At 8-weeks, the injured attachments had increased evidence of fibrovascular scar tissue in addition to fatty infiltration of the tendon compared to sham attachments (Figure 3D–D’). Collagen production/remodeling was apparent, indicated by the blue+ stain, at both 3- and 8-weeks after injury (Figure 3B’ and Figure 3D’).

**Figure 3.**
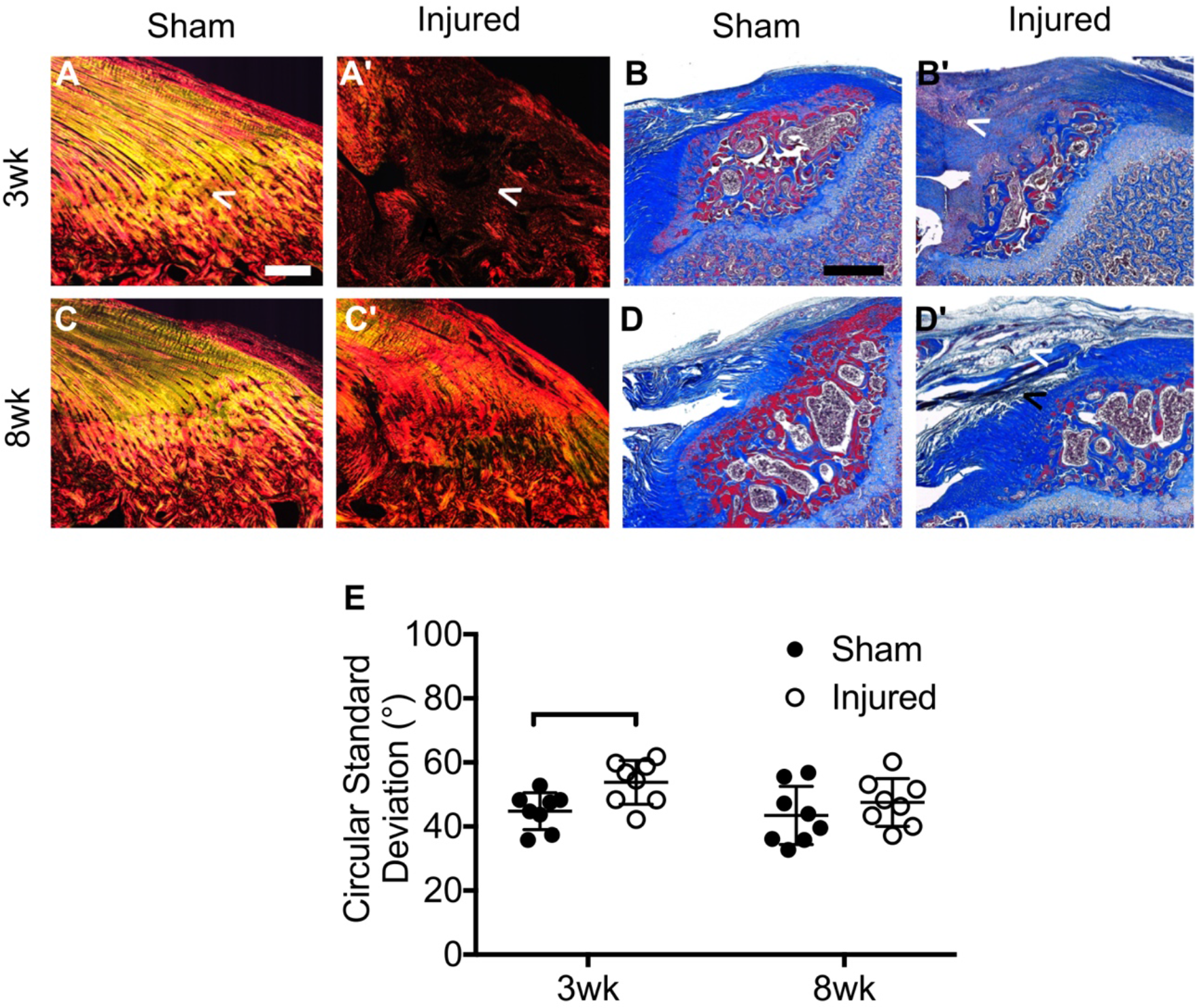
Organization at the IS attachment was impaired in 3wk injured attachments compared to 3wk sham attachments. (A-C’) Transverse histology stained with Picrosirius Red of (A) 3wk sham, (A’) 3wk injured, (C) 8wk sham, and (C’) 8wk injured. (B-D’) Transverse histology stained with Masson’s Trichrome of (B) 3wk sham, (B’) 3wk injured, (D) 8wk sham, and (D’) 8wk injured. White arrowheads in A & A’ highlight regions of organized collagen, with (A) sham attachments having noticeably more organized collagen compared to (A’) injured attachments. White arrowheads in B’ highlights ECM deposition at the injured IS attachment, as well as (D’) fatty accumulation in the tendon. Black arrowhead in (D’) highlights fibrosis at the injured attachment. (E) Circular Standard Deviation (quantification of collagen fibril alignment) was significantly increased for 3wk injured attachments compared to sham. Bars indicate significant differences between groups (p<0.05, mean ± 95% CI). Images taken at 5X magnification, scale bar: 200µm.

At 3-week post-injury, the attachments appeared unorganized and/or had non-existent organized collagen at the injury site (Figure 3A’). Sham attachments at both 3- and 8-weeks had organized collagen (yellow) at the tendon-bone attachment, with a clear transition in organization between tendon and bone (Figure 3A and Figure 3C). At 8-weeks post-injury, attachments demonstrated evidence of re-organization of collagen at the injury site compared to the 3-week injured attachments (Figure 3C’). The 3-week injured attachments had, quantitatively, decreased collagen organization compared to sham attachments (Figure 3E).

TRAP-positive osteoclasts were prevalent at the injured attachment, indicating an increase in bone remodeling at both 3- and 8-weeks (Figure 4A–D’). Injured attachments had increased Oc.S./B.S. at 3- and 8-weeks compared to the contralateral sham attachments (Figure 4F). Oc.S./B.S. was significantly higher in the injured 3-week attachments compared to the injured 8- week attachments (Figure 4F). There was no significant difference in Oc.S./B.S. for 3-week sham attachments compared to 8wk injured attachments. The overall cell density was significantly higher for both time points at the injured tendon-bone attachments compared to the sham attachments (Figure 4E).

**Figure 4.**
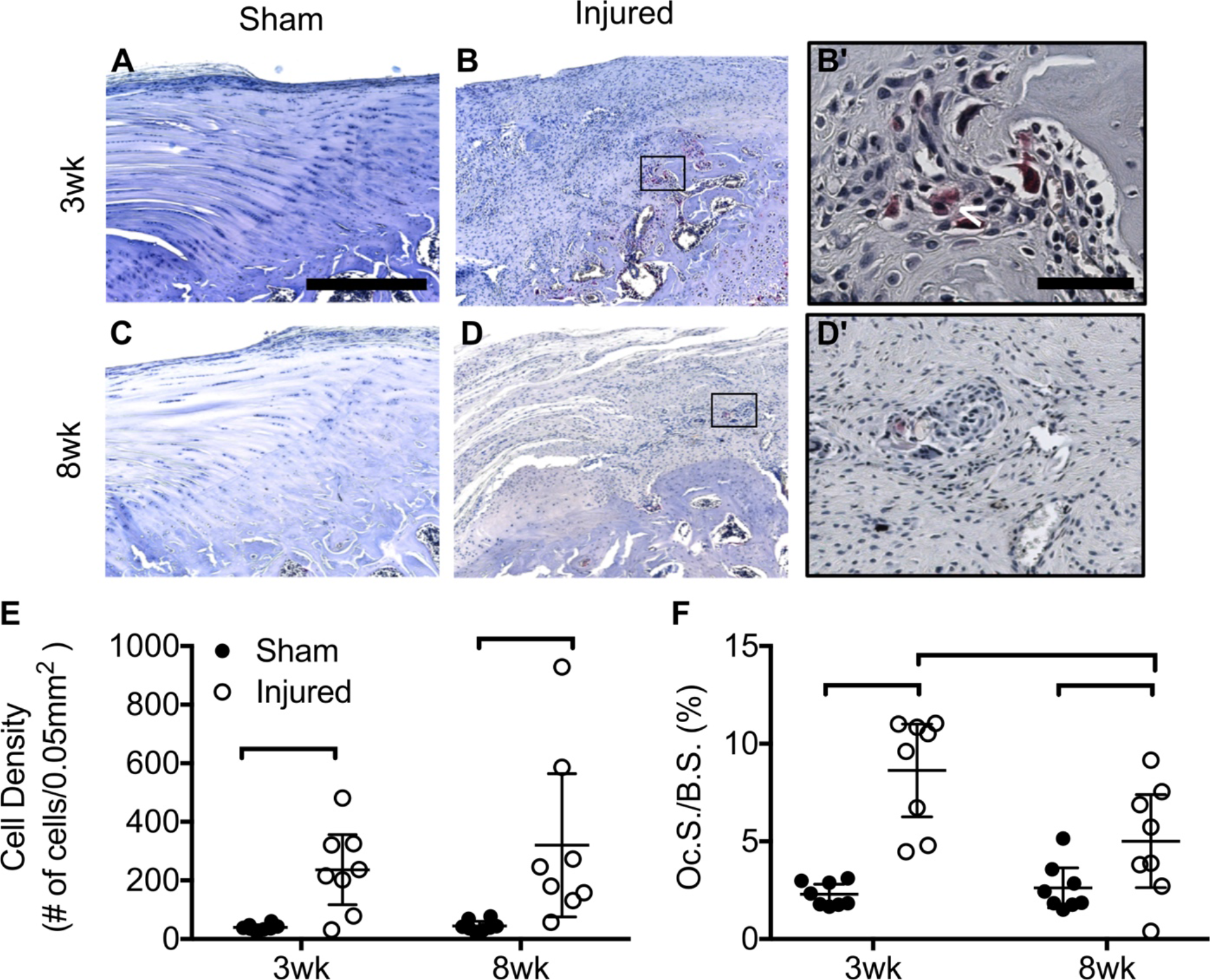
Cell Density and Oc.S./B.S. ratio was significantly greater in the injured attachments compared to the sham-operated attachments for both time points. (A-D) Transverse histological images at 5x-magnification of (A) 3wk sham, (B) 3wk injured, (C) 8wk sham, and (D) 8wk injured. Scale bar: 200µm. (B’-D’) 20x-magnification of (B’) 3wk injured and (D’) 8wk injured. Scale bar: 30µm. (E) Cell density and (F) Osteoclast surface (Oc.S.) to bone surface (B.S.) ratio. Bars indicate significant differences between groups (p < 0.05, mean ± 95% CI).

## DISCUSSION

### Overview

The structure of the healthy tendon-bone attachment is ideal for its function to transmit forces generated by muscle to bone (23, 25-27). Although the attachment resists failure, injuries commonly occur near the attachment within rotator cuff tendons (6, 17, 28-30). Rotator cuff tears may necessitate surgical repair of the tendon back to bone, however conservative treatments can also be effective in reestablishing cuff strength and health (2, 31-35). In the present study, we investigated cuff healing using a new model of partial width injury model in the rat.

Here, we used this newly-established injury model of a partial-width rotator cuff defect to quantitatively and qualitatively investigate the morphological properties of the healing attachment at 3- and 8-weeks post-injury.

### Study Significance

In this study, we showed that partial-width, full-thickness injuries led to substantial deficits in cellular and structural composition, including impaired collagen alignment, increased osteoclast activity, and decreased bone volume compared to sham-operated attachments. These data support that healing of the mature tendon-bone attachment is scar-mediated, coinciding with previous work (21). In our study, short-term following injury (3-weeks) led to decreased collagen organization compared to uninjured groups, yet the attachment was able to somewhat overcome this deficit in the longer term (i.e., by 8-weeks post injury). This suggests that the attachment continues to remodel and re-organize the injury site. Notably, the biopsy punch used to create the injury permeated the cortical bone, which may have resulted in increased osteoclast activity at the injury site, indicative of a bone-marrow derived healing response (21, 36). The observed increase in bone resorption, driven by osteoclast activity, likely led to reduced BV/TV in the injured groups at 3- and 8-weeks post-injury. Additionally, the injury may have led to reduced loading, due to pain or discomfort, resulting in a reduction in muscle forces transmitted to bone, which could have also contributed to the decreased BV/TV that we observed (8). Reduced muscle loading of the tendon-bone attachment is detrimental to tendon-bone healing (7, 8), yet it remains unknown if a partial-width injury to the tendon-bone attachment influences the muscle loading of the attachment in our model. Per our *ex vivo* validation studies, the injured attachments likely have decreased biomechanical properties at time of injury due to loss of structural integrity and removal of attachment tissue. Interestingly, none of the partial-width injuries resulted in a full-width injury, supporting recent evidence that the attachment may resist failure even when damaged, albeit less so than a healthy attachment (23). Therefore, although full regeneration of the structure of the tendon-bone attachment was not observed, the scar tissue formed at the injury-site may be mechanically sufficient to resist tear propagation.

### Research Significance

Although tear propagation was not observed in this study, rotator cuff tears in the clinic may propagate into full-width injuries that require surgical repair (37–39). Therefore, partial injuries to rotator cuff tendons are sometimes surgically repaired to prevent tear propagation (17, 28). Hence, many studies have aimed to understand tear propagation (23, 38–41) and various scenarios of rotator cuff repair (6, 17, 28–30, 33, 42–46). Despite the plethora of research in the field of rotator cuff repair, repairs often fail due to poor re-integration of tendon to bone (6, 47). As such, studies of attachment healing have focused on tissue-engineering approaches for improving attachment repair and establishing the mechanical and biological factors that are involved in repair healing (7–9, 14, 18, 19, 21, 22, 31–33, 36, 48–61). Few studies have investigated the innate healing response of the attachment (i.e., attachment healing without repair). One such study identifies a role of Gli1+ cells, key regulators of attachment development, in attachment regeneration via direct punch-like injury to neonatal mouse attachments (21). Although these cells mediate neonate regeneration, Gli1+ cells were scarce during scar-mediated healing in the mature attachment (21). Interestingly, scar-mediated healing has been shown to occur both in the mature mouse and in our rat model of healing, indicating that aging may limit the capabilities for tissue regeneration, regardless of species. The results of our study are consistent with other animal models of rotator cuff injury (7, 8, 10, 18, 19, 54, 62), elucidating that the composition of the attachment varies at different time points after an acute injury. In one study of acute rotator cuff tendon injury and repair, attachments exhibited a wound healing response with a lack of restoration achievable compared to uninjured shoulders (14). Only a few studies, however, have focused on the healing composition of partial-width injuries without repair or augmentation (19). In this study, we established that a model of partial-width rotator cuff injury in the rat shoulder leads to achievable restoration, although scar-like and fibrotic, at the attachment without surgical repair.

### Clinical Significance

An increased understanding of the healing properties of partial-width rotator cuff injuries are necessary to establish optimal treatment plans for patients with these injuries. The small-animal injury model presented here will be beneficial in determining the innate healing properties of the attachment, which can contribute to pharmacological interventions or treatment protocols for partial-width tears.

### Limitations

This clinical relevancy of this model is somewhat limited, as most partial-width tears reside on the bursal side of the attachment and include the lateral attachments of the tendon to bone (18, 32, 33). However, this model is useful to evaluate the injury and healing characteristics of the attachment, directly. Additionally, in this model, the punch biopsy permeated the cortical bone of the humeral head and removed a portion of bone, which could be contribute to the loss of BV/TV in the injured attachments compared to the sham operated attachments. Furthermore, the differences in time points and sex among the rats may contribute to slight variations in the healing of the attachments. The discrepancy between the BV/TV for the entire humeral head and injury region may be due to the sensitivity of the threshold parameters of the microCT software, resulting in undeveloped or less dense bone not being distinguished. Lastly, biomechanical testing was not performed for this injury *in vivo*. Thus, the strength of the integrity of the attachment is unknown, despite decreased structural quality at the injury site of the attachment after healing.

### Conclusions

A partial-width, full-thickness injury to the IS attachment of the rotator cuff compromises the quality of the attachment. This study established the structural quality of the attachment following healing in a new rodent model. Proliferation and remodeling of the attachment during healing resulted in attachments that were structurally inferior compared to an uninjured attachment even after 8-weeks of healing. This study contributes a novel *in vivo* injury of a partial-width, full-thickness rotator cuff injury that can be used as the basis for further research to evaluate healing properties and tear propagation of the attachment with moderate to vigorous exercise, immobilization, or pharmacological interventions.

## ACKNOWLEDGEMENTS

We acknowledge Terry Kokas, Histology Specialist III (Nemours-A.I. duPont Hospital for Children Histochemistry and Tissue Processing Core; supported by NIH-NIGMS: P20 GM103446 and the state of Delaware) and Crystal Idleburg, HT (ASCP) (Washington University Musculoskeletal Research Center, NIH NIAMS: P30 AR057235) for assistance with histology. The injury illustration (Figure 1A) was created by Katelyn McDonald, MA, CMI (Certified Medical Illustrator). Elahe Ganji assisted in creating the three-dimensional reconstructions of the injury model.

